# De Novo Computational Design of VHH Nanobodies Against LGR5

**DOI:** 10.64898/2025.12.02.691880

**Authors:** Chenrui Xu, Yifan Li, Trang Nguyen, Yi Zhou, Longfei Cong, Tek-Hyung Lee, Yuxiang Lang, Roger Shek, Li Yi, Per Greisen

## Abstract

VHH discovery traditionally relies on animal immunization or large-scale library screening, methods that are slow, costly, and often ineffective for challenging targets such as GPCRs. We present a fully de novo computational pipeline for epitope-directed VHH design, integrating generative backbone modeling, deep learning–based sequence optimization, and iterative experimental feedback. Using LGR5 as a model, we progressed from in silico design to functional binders without structural templates. Across three design–test–learn cycles, millions of candidates were reduced to epitope-specific binders with nanomolar affinity and high thermal stability (melting temperature [Tm] > 65 °C). Cryogenic electron microscopy (cryo-EM) confirmed atomic-level agreement (RMSD ≈ 2.2 Å). This structure-validated approach accelerates timelines, reduces cost, and is broadly applicable to GPCRs and other membrane proteins, enabling “on-demand” therapeutic antibody generation.

## Introduction

Antibodies have revolutionized modern medicine, with over 150 approved therapeutic antibodies generating more than $200 billion in annual revenue^1^. Their exquisite binding specificity, combined with diverse effector functions, has enabled breakthrough treatments for cancer, autoimmune diseases, and infectious diseases. However, conventional antibody discovery through animal immunization or library screening remains slow, expensive, and often fails for challenging targets such as membrane proteins, cryptic epitopes, or antigens requiring precise geometric constraints^2^. Single-domain antibodies (VHHs), derived from camelid heavy-chain-only antibodies, have emerged as a particularly promising class of biologics^3^. Their small size (∼15 kDa), exceptional stability, ability to access cryptic epitopes, and straightforward expression in microbial systems make them ideally suited for applications ranging from oncology to diagnostics^4^. Because VHHs consist of a single variable domain, they provide an ideal testbed for de novo computational design, offering a tractable system to explore direct sequence-to-function mapping.

The unique properties of VHHs—including their extended complementary-determining region (CDR) H3 loops capable of penetrating receptor cavities, high thermal stability (often retaining activity above 70 °C)^25^, and resistance to aggregation—have enabled targeting of previously undruggable epitopes^5^. Recent clinical successes, such as caplacizumab for thrombotic thrombocytopenic purpura^6^ and ongoing trials targeting tumor-associated antigens, demonstrate their clinical utility. Among these, membrane proteins such as leucine-rich repeat-containing G-protein coupled receptor 5 (LGR5)—a critical stem cell marker and emerging cancer target^7,8^—illustrate the discovery challenges that motivate this study. Current discovery methods—whether through camelid immunization, phage display, or synthetic library screening—require months of iterative selection, often yield low hit rates for difficult targets, and provide limited control over epitope specificity^9^. The inability to rationally design VHHs targeting specific epitopes remains a fundamental bottleneck in exploiting their full translational potential.

The past three years have witnessed remarkable progress in de novo protein design, driven by deep learning architectures such as RFdiffusion^10^, ProteinMPNN^11^, and AlphaFold^12^. These tools have enabled the design of protein binders, enzymes, and scaffolds with unprecedented accuracy, achieving experimental success rates exceeding 50% for some applications^13,14^. However, their application to functional antibody and VHH design remains limited. While these methods excel at generating stable protein structures, they struggle to simultaneously optimize binding specificity, affinity, and developability without extensive experimental validation^15^. Despite these advances, no integrated computational–experimental framework yet enables rapid, epitope-directed VHH generation with iterative feedback. The resulting gap between computational prediction and experimental validation means that designed molecules may be structurally sound yet functionally inadequate. Bridging this divide requires a unified design–build–test–learn pipeline that explicitly couples in silico prediction to high-throughput functional readouts^16^.

Here we present a fully de novo computational design pipeline for generating functional VHHs against specified epitopes without requiring structural templates, existing binders, or immunization. Our approach integrates state-of-the-art protein design algorithms with an experimental feedback loop using yeast surface display (YSD), enabling rapid optimization across multiple design cycles. We demonstrate this platform by targeting LGR5, designing VHHs with nanomolar affinity and epitope specificity validated through N-linked glycan scanning and competitive binding experiments. Structural determination by cryogenic electron microscopy (cryo-EM) confirms atomic-level accuracy of our designs. While exemplified here for LGR5, the same principles are broadly applicable to other membrane proteins and conformationally sensitive epitopes. Beyond providing functional LGR5 binders, this work establishes a generalizable framework for rapid, epitope-directed VHH discovery, with broad implications for therapeutic development and basic research targeting challenging membrane proteins and protein–protein interfaces.

## Results

We developed an iterative computational pipeline for designing VHH antibodies targeting preselected epitopes (**Fig. 1**). As a therapeutic target, we selected LGR5, a G-proteincoupled receptor (GPCR) with limited structural data regarding antibody or VHH interactions. This scarcity of structural information posed challenges for evaluating design accuracy, necessitating robust computational strategies.

**Figure 1.**
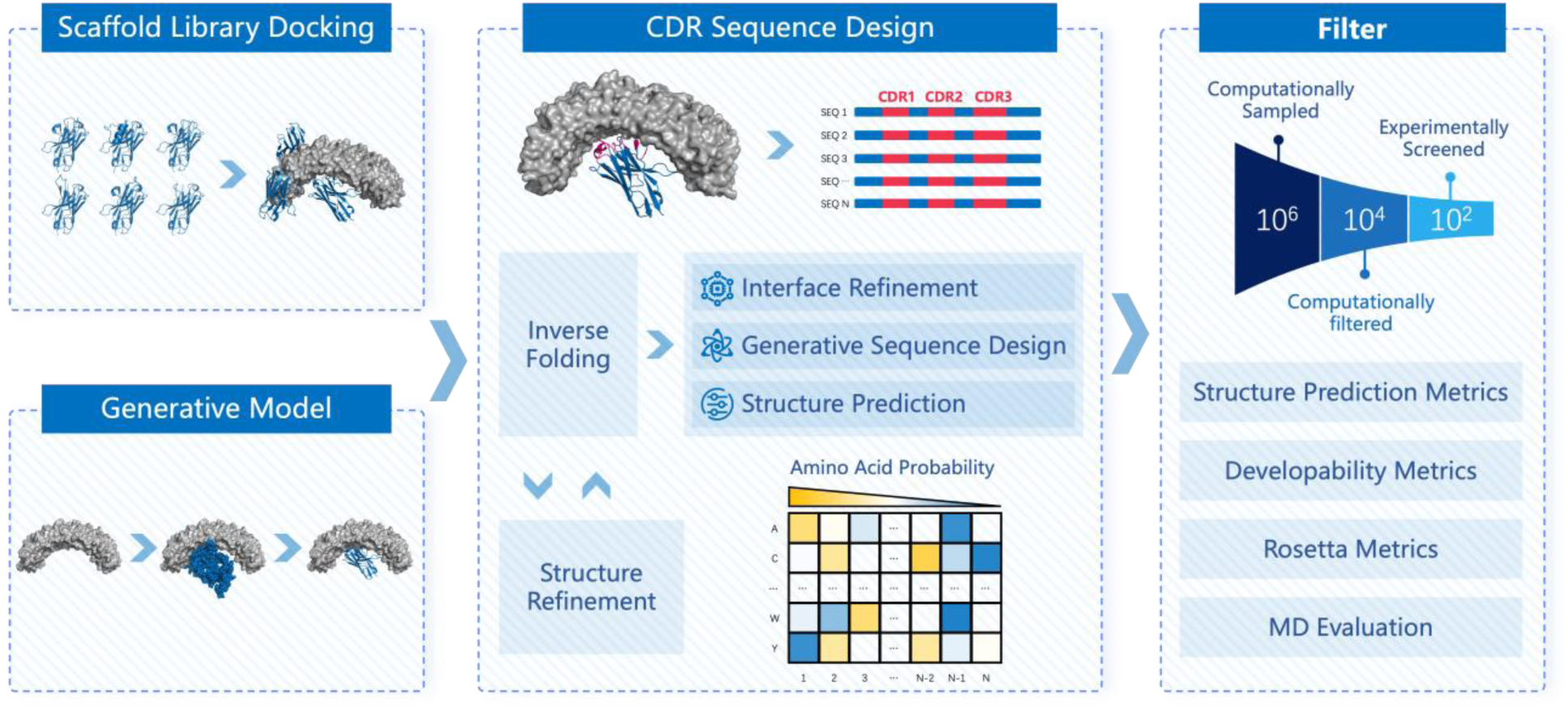
Computational workflow for de novo design of VHH. The pipeline integrates scaffold library docking and generative backbone modeling to create candidate structures. These feed into CDR sequence design, combining inverse folding, structure refinement, and generative sequence optimization. Designs are evaluated using structural and interface metrics, then filtered through computational scoring (structure prediction, developability, Rosetta, MD) to reduce millions of candidates to a small set for experimental screening.

### First-Round Design

In the initial stage, we developed and implemented the Epitope-Guided Generative Design (EGGD) Pipeline, a modular workflow that combines epitope-specific docking, generative backbone modeling (RFdiffusion), and deep learning–based sequence optimization (ByProt)^29^. This pipeline enables the de novo generation of VHH candidates tailored to specific antigenic surfaces, facilitating rapid exploration of sequence– structure–function relationships. To ensure well-defined and interpretable VHH–antigen interfaces, a hotspot-guided protein-protein interaction (PPI) generation strategy was constructed where the hotspot coordinates were derived from:

- Inverse rotamer sampling (e.g., RifGen)
- Existing ligand-bound complexes
- Manually curated models

The resulting scaffolds were docked to the target antigen using an in-house docking protocol. This protocol leverages predefined hotspot interaction pairs to determine the optimal relative orientation between the VHH scaffold and the antigen. Each docking pose was evaluated based on three key criteria: (i) proper engagement of the designated hotspot residues on the VHH, (ii) minimal root-mean-square deviation (RMSD) between the positioned residues and their original locations on the scaffold, and (iii) absence of significant steric clashes between the antigen and the VHH framework. After the initial placement of the hotspot residues, we performed CDR loop and interface optimization using a multi-step protocol that combines backbone sampling (RFdiffusion), sequence generation (ByProt), and physics-based refinement (Rosetta). To enhance the quality and success rate of candidate designs, we applied a rigorous filtering strategy based on structural and energetic metrics derived from experimentally validated antibody–antigen complexes in the Protein Data Bank (PDB). Key metrics included shape complementarity (SC), buried solvent-accessible surface area (ΔSASA), and the interface energy normalized by surface burial (ΔG/ΔSASA ratio). These parameters were selected for their strong discriminative power in identifying structurally and functionally viable binders, thereby reducing the candidate pool while enriching high-confidence designs.

Fluorescence-activated cell sorting (FACS) analysis of the initial design library demonstrated that over 60% of the designs were robustly displayed on the surface, suggesting proper protein folding and structural integrity during preliminary evaluation^30^. However, the antigen-based binding screening only identified 6 potential binders with an estimated affinity of more than 500 nM. These 6 binders did not pass the FACS epitope specificity characterization using N-linked glycan LGR5 variants and were not advanced to biophysical characterization (**Table** 1). To address the limited success rate, we incorporated AlphaFold2 (AF2)-based interface redesign and expanded scaffold diversity in the next round.

**Table 1.** Summary of Design–Validation Iterations.

| Round | Designs Tested | Passed Display (Folding/Foldeness) | Passed Binding (FACS-Based analysis) | Passed Epitope Specificity (FACS-Based analysis) | Advanced to Biophysical |
| --- | --- | --- | --- | --- | --- |
| 1 | 2576 | >1500 | 6 | NA | NA |
| 2 | 2000 | >1500 | >30 | 5 | 2 |
| 3 | 35 | 28 | 7 | 7 | 3 |

### Second-Round Design

First-round designs showed low hit rates and structural inconsistencies when evaluated by folding models such as AF2, with altered epitopes or reduced interface predicted Template Modeling (ipTM) and interface predicted Local Distance Difference Test (ipLDDT) scores. To address these limitations and improve both structural fidelity and functional success, the pipeline was further optimized by integrating:

- Pose sampling to generate diverse initial antigen–VHH orientations
- AF2-based redesign of interface residues
- Sequence optimization of non-interface CDRs
- Structural self-consistency filtering following re-folding

Specifically, a comprehensive VHH scaffold library was constructed from the PDB, VHH sequence repository (Integrated Nanobody Database for Immunoinformatics (INDI) comprising patent- and next-generation sequencing (NGS)-derived sequences ^31^) to expand the scaffold library. For VHHs without experimentally solved structures, we generated apo VHH models using xTrimoFast^32^ and incorporated only high-confidence predictions (pLDDT > 90) into the scaffold library^32^. Initial poses were generated using AF2-Multimer and geometric docking, then refined through AF2-based interface redesign. Non-interface CDRs were optimized with ByProt, and thousands of candidate complexes were filtered using multi-criteria scoring like Boltz2 and Chai-1 to test for structural self-consistency.

Like the first-round constructs, most second-round designs (>75%) exhibited efficient surface expression on yeast cells, qualifying them for subsequent antigen-binding and epitope characterization. More than 30 designs were identified to be effective LGR5 binders, with estimated affinity ranging from 100 nM to >1 µM. FACS-based analysis indicated that 5 designs might target the designed epitopes, among which 2 designs were further advanced for biophysical characterization. However, those designs did not undergo detailed characterization, including competitive binding and complex structure determination, either due to low affinity or poor biophysical properties (**Table** 1).

### Third-Round Design

Second-round designs showed improved folding consistency and higher yeast display success, but computational models such as Boltz-2 continued to favor epitopes near the natural ligand-binding site, limiting epitope diversity. Epitopes associated with native ligand binding yielded higher confidence scores, causing uneven distribution of targeted epitopes. The top 10% designs with the highest re-folding consistency scores were targeting the same epitope as the natural ligand. To mitigate this, we modified the Boltz2 pipeline to incorporate the original design complex as a structural template, constraining the model toward the intended docking geometry. This in silico adjustment improved the concordance between the initially intended epitope—defined by the design input—and the epitope predicted after sequence optimization and modeling of key interactions, as validated through self-consistency with the structure prediction algorithm. Further analysis revealed two issues with ipTM-based filtering:

- Enrichment of hydrophobic residues in designed regions.
- Insufficient prioritization of specific interfacial interactions critical for functional binding.

To reduce exposed hydrophobic residues in the CDRs, we required that hydrophobic residues be involved in molecular interactions: π–π, cation–π, and charge-polar interactions, which were identified via geometric contact analysis^24^.

### Outcome

After three rounds of computational refinement and experimental feedback, we prioritized three lead VHH designs out of seven identified variants for detailed experimental validation (**Table 1, Fig. 2**). These candidates demonstrated consistent display efficiency, epitope-specific binding on YSD, and favorable computational metrics, making them suitable for biophysical characterization and structural analysis.

**Figure 2.**
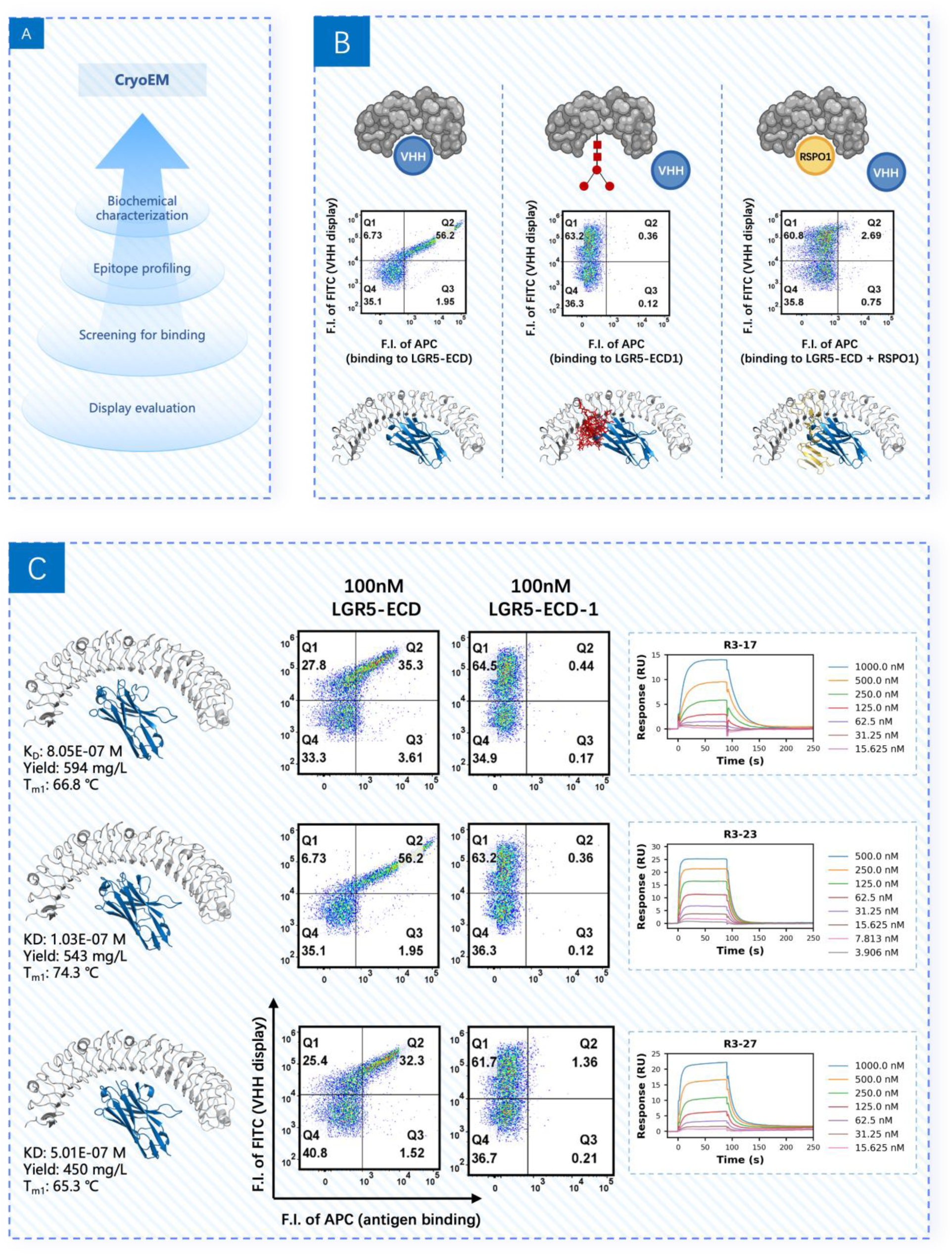
Experimental screening and characterization of designed VHHs. (A) Workflow for experimental evaluation of computationally designed VHHs, including display evaluation, binding screening, epitope profiling, and structural validation by cryo-EM. (B) Epitope mapping of selected VHHs on LGR5 ECD. Flow cytometry plots show binding profiles for different epitope classes, with corresponding structural models highlighting binding regions on LGR5. (C) Biophysical characterization of lead VHH candidates. Binding affinity measurements by SPR are shown alongside flow cytometry binding curves. Representative cryo-EM models of VHH–LGR5 complexes are displayed for each candidate.

### Final Validation and Structural Confirmation

To rigorously validate the computational pipeline and establish a data-driven optimization framework, we developed a high-throughput screening platform leveraging yeast surface display (YSD) (**Fig. 2**). This system enables rapid assessment of folding integrity and antigen-binding properties of *in silico* designs. To enforce epitope-specific recognition rapidly rather than nonspecific interactions, we incorporated an N-linked glycan masking strategy during selection, both during library sorting and single clone characterization^17^. This approach designs computationally N-linked glycans on the target epitope, effectively occluding the designed interface and providing a stringent filter for specificity.

### FACS-based analysis of in silico designs

The experimental workflow began with cloning computationally designed variants into YSD constructs, followed by evaluation of surface display efficiency. YSD serves as a proxy for correct folding^19^, as higher display levels generally correlate with improved structural integrity during recombinant expression^18^. From the third design iteration, 35 candidates were screened. Of these, 28 exhibited robust display signals and were advanced to antigen-binding analysis.

Binding assays were performed using FACS against 100 nM LGR5 extracellular domain (LGR5-ECD). Seven designs demonstrated clear and reproducible binding signals. To confirm epitope specificity, these candidates were investigated with N-linked glycan LGR5 variants (LGR5-ECD-1 and LGR5-ECD-2) as well as two natural competitors, R-spondin 1 (RSPO1) and R-spondin 3 (RSPO3). This dual approach was chosen because the natural ligand covers a larger area including the intended epitope while the N-linked glycans mask the intended epitope at near-atomistic resolution. The masking strategy was validated using YSD RSPO1^17^, which bound native LGR5-ECD but failed to interact with N-linked glycan variants at 100 nM^26^, confirming effective occlusion of the targeted epitope (**Suppl. Fig. S1**).

All seven designs exhibited a consistent binding profile: strong interaction with unmodified LGR5-ECD, but near-complete loss of binding to glycan-masked variants at 100 nM. Furthermore, competitive binding assays revealed that the addition of 250 nM RSPO1 or RSPO3 abolished interaction with LGR5-ECD^22^, indicating that the designed binders target the RSPO-binding epitope on LGR5 in correspondence with the computational model.

### Biophysical characterization of selected in silico designs

Six of the seven designs (POC2-R3-03, -17, -19, -23, -24, and -27) identified through FACS pre-characterization were advanced for recombinant production and *in vitro* biophysical analysis. The recombinant production and purification results showed that Fc-fused designs were all produced with POC2-R3-17, -23, and -27 being successfully purified in single valent VHH-6xHis format. These three binders were selected for detailed affinity and Tm determination. The surface plasmon resonance (SPR) results confirmed the binding of all 3 binders against LGR5-ECD with affinity (dissociation constant, K_D_) in the range of 100 nM ∼ 1 µM. It is worth noting that melting temperature (Tm) measurements of all three designs exceeded 65 ℃, indicating good thermal stability of these designs. Finally, POC2-R3-23, as the top binder with a K_D_ of approximately 100 nM, was selected to be structurally validated by Cryo-EM.

### Structural Validation of VHH-LGR5 Complex

To maximize the quality of the structure, the VHH-LGR5-ECD complex was prepared with an additional non-competing fragment antigen-binding (Fab) to increase the molecular weight thereby enhancing the signal-to-noise ratio (SNR) during data collection. Only one single binding pose was observed during 3D classification (**Suppl. Fig. S2**), suggesting high specificity in binding to the designed epitope. During 2D classification, diverse orientation projections were observed, among which our designed VHH exhibited a strong density at the intended epitope (**Fig. 3A**). The Cryo-EM structure had an overall resolution of 3.19 Å, with high confidence side-chain placement at the VHH-Ag interface except for the CDR3 region. The atomic complex structure was constructed using this high-quality density map. The experimentally determined apo-VHH shows excellent agreement with our computational model, exhibiting an overall RMSD of 0.5 Å, with local RMSDs for each CDR all below 2 Å (**Fig. 3B and 3C**). The high-resolution holo-complex showed structural agreement at the interface region. The VHH orientation aligned well between the computational model and the Cryo-EM structure, and the hotspot interacting residues coincided with atomic-level precision (**Suppl. Fig. S2**).

**Figure 3.**
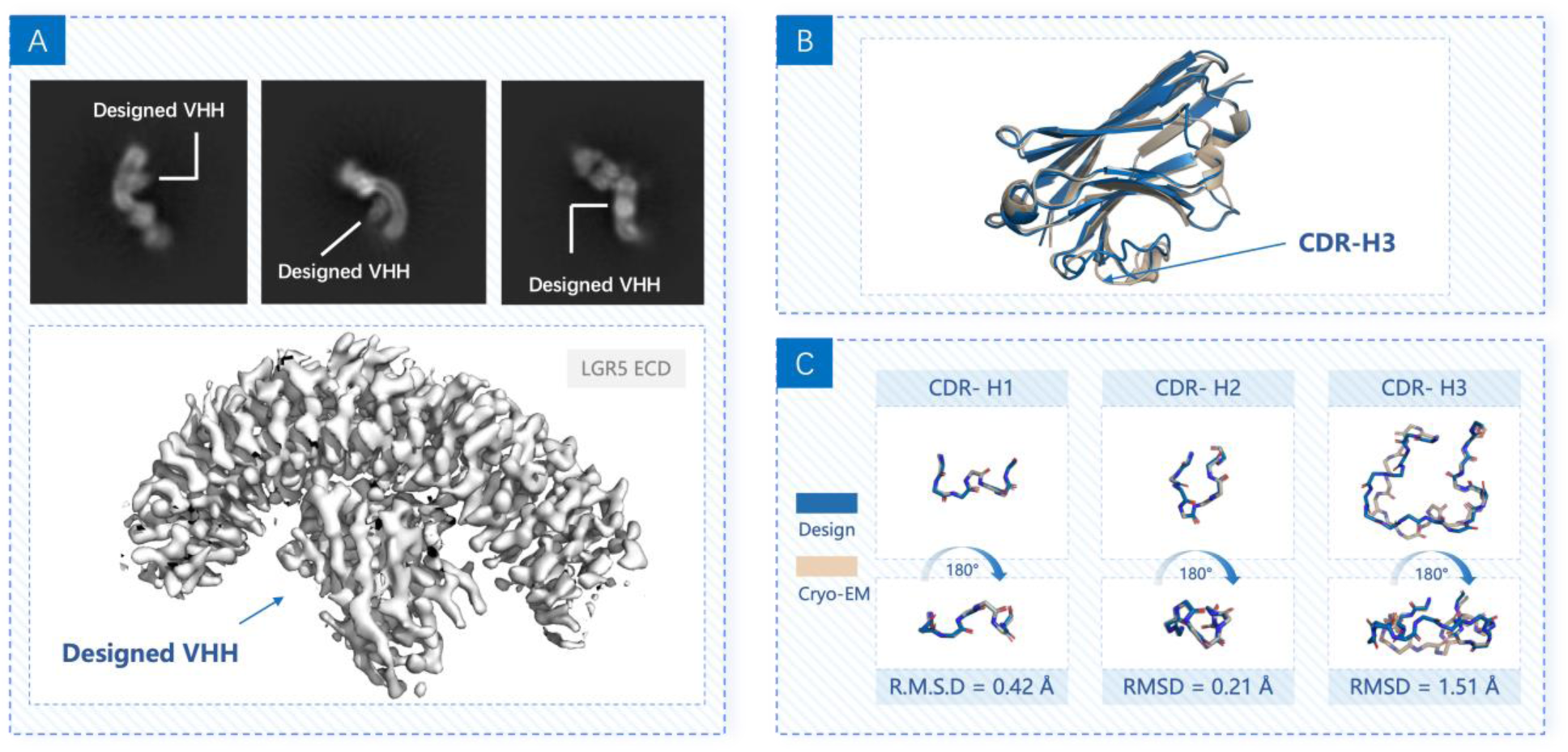
Structural characterization of designed VHH bound to LGR5 ECD. (A) Representative 2D class averages of cryo-EM particles showing the designed VHH in different orientations (top) and the cryo-EM density map of the LGR5 extracellular domain (ECD) in complex with the designed VHH (bottom). The VHH density is indicated by arrows. (B) Structural overlay of the designed VHH model (tan) and cryo-EM structure (blue), highlighting the CDR-H3 loop. (C) CDRs between design and cryo-EM structure for CDR-H1, CDR-H2, and CDR-H3. RMSD values are shown for each loop, with CDR-H3 exhibiting the largest deviation (1.51 Å).

## Discussion

We report the demonstration of fully de novo computational design of VHH nanobodies against a structurally challenging membrane protein target, LGR5. By integrating generative backbone modeling, epitope-guided docking, and deep learning–based sequence optimization with iterative experimental feedback, we established a design– build–test–learn pipeline that bridges the gap between structural prediction and functional validation. Across three design cycles, this approach progressed from modest initial hit rates to epitope-specific binders validated by cryo-EM at near-atomic resolution, underscoring the feasibility of rational antibody design without reliance on immunization or pre-existing templates.

Compared to conventional discovery strategies—camelid immunization, phage display, or synthetic library screening—our pipeline offers distinct advantages in speed, cost, and epitope control. Traditional workflows require months of iterative selection and often fail for GPCRs and other conformationally sensitive targets. In contrast, our computational framework generated nanomolar-affinity binders within weeks, guided by structural and physicochemical filters that prioritize folding integrity, interface complementarity, and developability. Although early designs exhibited limited affinity and specificity, successive rounds incorporating AlphaFold-based interface redesign and scaffold diversification markedly improved success rates, highlighting the importance of coupling in silico prediction with experimental screening.

The implications of this work extend beyond LGR5. GPCRs and other membrane proteins remain among the most challenging classes for antibody discovery, and the ability to rationally design VHHs targeting predefined epitopes enables “on-demand” generation of therapeutic candidates. This capability could accelerate timelines for oncology, immunology, and diagnostic applications, and facilitate rapid responses to emerging pathogens or personalized medicine needs. Integration of this pipeline with downstream therapeutic development frameworks—including affinity maturation, Fc engineering, and manufacturability assessment—positions computational design as a transformative tool for biologics discovery.

Future directions include expanding the platform to design multi-specific binders, optimizing constructs for therapeutic modalities such as bispecific antibodies or chimeric antigen receptor T-cell (CAR-T) components, and extending the approach to other antibody formats beyond VHHs. Incorporating higher-throughput biophysical screening and structure-guided optimization strategies will further enhance affinity and stability. Ultimately, coupling this workflow with in vivo functional assays and developability profiling will enable seamless progression from computational design to clinical candidates.

In summary, this study establishes a generalizable paradigm for computational antibody design, demonstrating that integration of generative modeling, predictive filtering, and experimental feedback can yield epitope-specific binders with atomic-level accuracy. By overcoming long-standing limitations of conventional discovery methods, this approach provides a blueprint for next-generation biologics development—faster, more precise, and adaptable to diverse therapeutic challenges.

## Methods

### Inverse-rotamer–based docking

Once inverse rotamers were defined, the customized docking algorithm performed exhaustive searches for compatible sites on VHH CDRs capable of accommodating hotspot rotamers. Compatibility between inverse rotamers and VHH residues was quantified using Cα and N atom coordinates for alignment. Candidate poses with severe steric clashes or insufficient CDR contacts were discarded, and favorable complexes were retained as templates for downstream optimization.

### AF2-Multimer sampling

AF2-Multimer predictions were run with five recycles, with apo structural templates, and model ensemble averaging over three independent seeds to enhance sampling diversity. Chain order was defined as (antigen–VHH), and inter-chain pairing restraints were disabled to allow unbiased complex formation. Predictions were ranked based on the ipTM and interface pLDDT scores. Among the sampled complexes, those exhibiting the desired epitope were chosen for further design optimization.

### AFdesign setup

The AFdesign binder protocol in ColabDesign was employed to refine the VHH CDR sequences^23^. Using the complex structure obtained from the previous step, the CDR regions were redesigned through AF2-based hallucination. Additionally, a position-specific scoring matrix (PSSM) derived from a protein language model was incorporated to bias the optimization toward functional and expressible molecules.

### RFdiffusion–ByProt–FastRelax pipeline

Building upon the **ProteinMPNN–FastRelax** paradigm, we developed a **RFdiffusion– ByProt–FastRelax** workflow that integrates deep generative modeling with physics-based refinement. For each VHH–target complex, RFdiffusion sampled alternative CDR conformations compatible with the antigen surface, while ByProt generated corresponding sequences that fold to the sampled structures. The combined models were threaded and relaxed with Rosetta **FastDesign/FastRelax**, wherein target backbones were fixed, and antibody backbones allowed flexibility. Sidechains of both partners were repacked for subsequent filtering.

### RFDiffusion for CDR sampling

CDR loops were sampled via partial diffusion with fixed framework and antigen coordinates (partial_T = 5, 10, 15). CDR loop regions were defined by the **Chothia** scheme^21^, and 20 designs were generated per complex using the **Complex_base_ckpt** model. The generated backbones replaced corresponding atoms in the template complex and were used for subsequent inverse folding.

### ByProt for CDR sequence generation

Based on the redesigned CDR coordinates from RFDiffusion, the **ByProt** inverse folding model generated sequence variants conditioned on fixed framework and antigen residues. Using a sampling temperature of 0.7, five sequences per structure were produced, yielding 100 total designs per docking pose.

### FastRelax protocol

All 100 designs underwent **FastRelax** optimization. During relaxation, antigen and noninterface residues remained fixed, while interface residues (within 8 Å) were repacked.

Interface residues received increased weighting (factor = 3) in the ref2015 score function. Metrics such as **sap_score**, **num_clashes**, and **contact_molecular_surface** were recorded.

### Expansion of the VHH Scaffold Library

To enhance structural diversity, VHH sequences were aggregated from **INDI**^31^ and clustered. Representative sequences were predicted with **xTrimoFast**^32^, and only models with ≥90% residues having pLDDT > 90 were retained, expanding the scaffold library from ∼700 to ∼7,000 entries.

### VHH Interface Design with AFdesign

Each initial pose underwent **AFdesign**-based optimization. Sequences with interface Predicted Aligned Error (iPAE) < 0.35 were used to construct **CDR-specific PSSMs** for targeted resampling. Validated sequences maintaining docking geometry and high AF2 confidence were advanced.

### Design Refinement with LLM and ByProt

We used a large language model (LLM) and **ByProt** to refine non-interface CDRs. Residues within 8 Å were defined as interfacial regions. Variants preserving epitope engagement and structural confidence in AF2 were retained.

### Interface SASA Ratio

This filter ensures that the designed VHH has sufficient contact with the antigen through the CDR or CDR3 regions. The ΔSASA for the entire VHH and each CDR region was calculated as the difference in SASA between the VHH in its apo and holo states. The detailed calculation procedure can be found in [https://github.com/biomap-research/DenovoVHH], and the distribution of ΔSASA-related metrics is presented in [https://github.com/biomap-research/De-novoVHH].

We observed that ΔSASA of CDR3 increases with longer CDR3 lengths ([https://github.com/biomap-research/De-novoVHH] for details). Using CDR3 ΔSASA as a filter would bias the design process toward VHHs with longer CDR3 regions. Therefore, we adopted the cdr_int_ratio as a filter, defined as the ratio of CDR ΔSASA to CDR SASA in the apo format of the VHH. As shown in [https://github.com/biomap-research/De-novoVHH], cdr_int_ratio exhibits almost no correlation with CDR3 length, making it a more robust metric for this purpose. We used closed intervals defined by 0.25–0.95 quantile to crystal ΔSASA. The cutoffs are:

- cdr3_interface_sasa_ratio between 0.3188 and 0.6965
- cdr_interface_sasa_ratio between 0.2862 and 0.535

### Shape Complementarity

The sc_value for all structures in the Structural Antibody Database (SAbDab)^33^ dataset and the designed models was calculated using the Rosetta InterfaceAnalyzer. We found that relaxation protocols influenced sc_value scores ([https://github.com/biomapresearch/De-novoVHH] for details). To align with our pipeline, we standardized the use of sc_value scores obtained after applying the FastRelax protocol.

### Energy-Related Scores from Rosetta

All energy-related scores were calculated using the XML protocol described in [https://github.com/biomap-research/De-novoVHH]. To identify which scores best differentiate SAbDab crystal structures from designed sequences, we created a dummy design dataset and compared the statistical distributions of all energy-related scores between the two groups ([https://github.com/biomap-research/De-novoVHH]).

We found that ΔG_separated/ΔSASA × 100, as well as the combined scores of side1_normalized + side2_normalized, provided the best distinction between the two groups. Consequently, these metrics were selected as filters for the design process.

### Sequence Optimization of Structural and Sequence Liabilities

For each potential pathological residue identified in a design, we first checked if it participated in any π-stacking or cation–π interactions with the antigen. If an interaction was found, we excluded the position from further mutations. Next, we checked if the closest germline residue was highly conserved. If the ImMunoGeneTics information system (IMGT) germline frequency of the germline residue was higher than 0.9, and the frequency of the current residue was lower than 0.05, the germline residue was considered conserved and limited as the only possible residue to be mutated to. Otherwise, all residue types in the mutational space for that position were included.

All positions that could be mutated were specified in a Rosetta Resfile^20^ with action “PIKAA” for mutating and repacking. VHH and antigen residues in contact with each other were tasked with action “NATAA”, meaning that they can be repacked but not mutated. All other residues were kept fixed with their native rotamers ([20]). We generated 30 optimized variants of each input design.

### Identification of Potential Pathological Residues

Complex structures were relaxed using Rosetta with all heavy atoms restrained to input coordinates. The “beta_nov16” score function was used with the “cart_bonded” term reweighted to 1.5, and “coordinate_constraint” reweighted to 1.0. All interatomic interactions in the relaxed structures were scanned with Arpeggia, a geometry-based interaction search tool we developed internally^27^. The following algorithms were used to identify residues with potential pathological behaviors:

- **Exposed hydrophobic residues:** all phenylalanine, tyrosine, tryptophan, leucine, isoleucine, and methionine with solvent accessible surface area (SASA) greater than 10 Å² and fewer than three hydrophobic contacts to other residues.
- **Buried polar residues without stabilizing interactions:** all serine, threonine, asparagine, glutamine, aspartic acid, glutamic acid, arginine, lysine, and histidine with (i) SASA less than 20 Å², (ii) no ionic bonding, hydrogen bonding, or polar contacts to other residues, and (iii) more than two hydrophobic contacts or van der Waals contacts to other residues.
- **Unfavorable ionic interactions:** any residue satisfying the “IonicRepulsion” term.

Sequence liabilities were identified via regular expression searches:

- N-linked glycosylation sites: “(N)[^P][ST]”.
- Asparagine deamidation sites: “(N)[GNHST]”.
- Aspartic acid sites: “(D)[GDHST]”.
- Methionine oxidation sites: “(M)”.

Filtering thresholds were determined by analyzing distributions from **SAbDab** VHH– antigen complexes and comparing crystal structures to RFDiffusion-generated decoy models, which served as negative samples. Metrics showing strong discriminative power between experimentally determined and generative complexes were retained, and the 15th percentile of the crystal-structure-derived distribution was used as the lower cutoff:

- CDR3 interface SASA ratio > 0.438
- Total CDR interface SASA ratio > 0.34
- Shape complementarity (Sc) > 0.6
- ΔG_separated / ΔSASA × 100 < –1.65
- side1_normalized + side2_normalized < –2.32
- spatial aggregation propensity (SAP) score between 40 and 70

### Filtering metrics with Boltz2 and Chai-1

Empirical thresholds for **Boltz2** and **Chai-1** were derived by re-folding post-2023 antibody–antigen crystal complexes. Outlier complexes with minimal interfacial contacts or excessively flexible loops were excluded from analysis. To identify metrics with the strongest discriminative power, we compared the score distributions of crystal structures and failed designs (all round-1 designs were used as negative examples). Scores that best distinguished crystal structures from failed designs were selected, and the median values of these metrics from the curated crystal dataset were adopted as thresholds. The following cutoffs were adopted:

- Boltz_iPTM > 0.75
- Boltz_iPLDDT > 0.90
- Chai_iPTM > 0.25
- H3_pLDDT > 0.84

### Sequence clustering and selection

Sequence clustering was performed using two complementary approaches. In the earlier rounds, we computed **xTrimo**-based sequence embeddings and applied t-distributed stochastic neighbor embedding (t-SNE) clustering to group sequences according to embedding similarity. From each cluster, the design exhibiting the lowest RMSD, highest H3_pLDDT, and highest Boltz_iPTM was selected. However, computing xTrimo embeddings proved computationally expensive for large sequence sets. Therefore, in later rounds, we adopted a more scalable strategy using **hierarchical agglomerative clustering** based on pairwise Levenshtein (edit) distances. A condensed distance matrix was first constructed from all pairwise sequence distances, providing a direct measure of sequence dissimilarity at the character level. The matrix was then subjected to average-linkage hierarchical clustering, iteratively merging the most similar clusters until a dendrogram was formed. Finally, the dendrogram was cut to yield a predefined number of clusters, allowing efficient partitioning of large design sets into representative, nonredundant groups without relying on alignment-based heuristics.

### Yeast surface display construct generation

35 designed VHH gene fragments were synthesized as gBlocks Gene Fragments Integrated DNA Technologies (IDT). Each fragment was cloned into the HindIII/BamHI-digested pXAGA2 vector using In-Fusion Cloning (Takara Bio) and transformed into *E. coli* NEB 5-alpha competent cells (New England Biolabs). Positive clones were identified by colony PCR, and purified PCR products were submitted to Genewiz for sequence verification. Sequence-confirmed constructs were subsequently transformed into Saccharomyces cerevisiae EBY100 cells using the Frozen-EZ Yeast Transformation II kit (Zymo Research) and plated on SDCAA agar plates for 2–3 days at 30 °C.

### FACS-based LGR5 binding and epitope characterization

For expression, a single colony was inoculated into SD-UT medium with glucose and grown overnight at 30 °C. Expression was induced by switching to SD-UT medium containing galactose and incubating at 25 °C for overnight. Approximately 2 × 10⁵ cells were collected per screening reaction, washed twice with phosphate-buffered saline (PBS) containing 2% bovine serum albumin (BSA), and labeled with Anti-FLAG Alexa Fluor 488 (AF488) antibody for 15 min at 4 °C followed by 15 min at room temperature. After washing, cells were incubated with the test antigen (either biotinylated or 6xHis tagged) at 37 °C for 30 min, then labeled with streptavidin–allophycocyanin (Avi-APC) for 10 min or anti-6xHis-APC antibody for 30 min at 4 °C prior to analysis using a NovoCyte flow cytometer. The antigen binding affinity was evaluated by the mean fluorescence intensity ratio of APC signals (antigen binding) over AF 488 signals (binder display on cell surface).

### Protein production and biochemical characterization

Each design DNA (codon optimized) was synthesized and cloned into pcDNA3.4, fused with a C-terminal hIgG1 Fc or 6×His tag connected by a 2(G4S) linker. Free cysteine in the hinge region was mutated to serine to avoid aggregation caused by disulfide bond mispairing. Expression was performed in ExpiCHO-S cells, and purification was carried out using either Protein A resin or Ni^2+^-NTA resin and further polished by SEC. VHH-Fc was eluted and stored in 0.1M HAc,130mM Tris pH5.5 and VHH-6xHis final product was stored in PBS. Purity of the products was evaluated by size-exclusion chromatography– high-performance liquid chromatography (SEC-HPLC) and capillary electrophoresis (CE).

### Antigen production and characterization

Human LGR5 extracellular domain (LGR5-ECD, residues 22–556, UniProt ID: O75473-1) was cloned into pcDNA3.4, fused with a N-terminal IL10 signal peptide (MHSSALLCCLVLLTGVRA) and a C-terminal Avi-6xHis tag (GGGGSGGGGSGGGGSGLNDIFEAQKIEWHEGHHHHHH). LGR5-ECD was expressed in ExpiCHO-S cells, followed by purification using Ni2+-NTA resin and further polished by SEC using Superdex 200 column on the ÄKTA protein purification system (GE Healthcare). Biotinylated LGR5-ECD was generated by co-transfecting tag-free BirA expressing plasmids at a molar ratio of BirA:LGR5 as 1:9. Purification of biotinylated LGR5 is the same as non-biotinylated LGR5, as described above. Final products were stored in PBS solution. Protein purity was evaluated by sodium dodecyl sulfate– polyacrylamide gel electrophoresis (SDS-PAGE), SEC-HPLC and oligomerization were evaluated by size-exclusion chromatography–multi-angle light scattering (SEC-MALS). Mass spectrometry post-translational modification analysis (MS-PTM) was performed to verify natural modifications are preserved by the CHO expression system.

### Bio-Layer Interferometry (BLI) Binding and Surface Plasmon Resonance Experiments

All BLI experiments used PBS with 0.02% Tween 20 and performed at 37℃.

For binder screening, 10 µg/mL VHH-Fc was captured using AHC2 Biosensor (Sartorius) with a target min of 1.0 response units. For screening, 1 µM LGR5-ECD (wildtype or mutant) was tested as analyte in single cycle kinetics with 240 s association and 300 s dissociation at rotation speed of 1000 rpm. After subtracting the reference signal from the sample signal, the binding kinetics were calculated using ForteBio data analysis software (version 12.0), and a 1:1 binding model was applied for curve fitting of LGR5 binding to VHH-Fc.

The binding affinity of biotinylated-LGR5-ECD to VHH-6xHis was determined using a Biacore 8K (Cytiva) instrument and Series S Sensor Chip SA (Cytiva). 1×HEPES (10 mM HEPES, 150 mM NaCl, 3 mM ethylenediaminetetraacetic acid (EDTA) with 0.005%Tween-20; pH7.4 was used as the running buffer. 5 μg/mL biotinylated-LGR5ECD was immobilized on SA Chip with a target immobilized level of 400 response units at a flow rate of 10 μL/min. 2-fold dilutions of each nanobody were tested in multi-cycle kinetics with 90 s association and 210 s dissociation at a flow rate of 30μl/min. SPR signals were acquired and analyzed using Biacore Insight Evaluation Software and fitted by 1:1 binding or Steady state affinity model to determine the K_D_.

## Supplementary Results

### Supplementary Tables

**Table S1.**
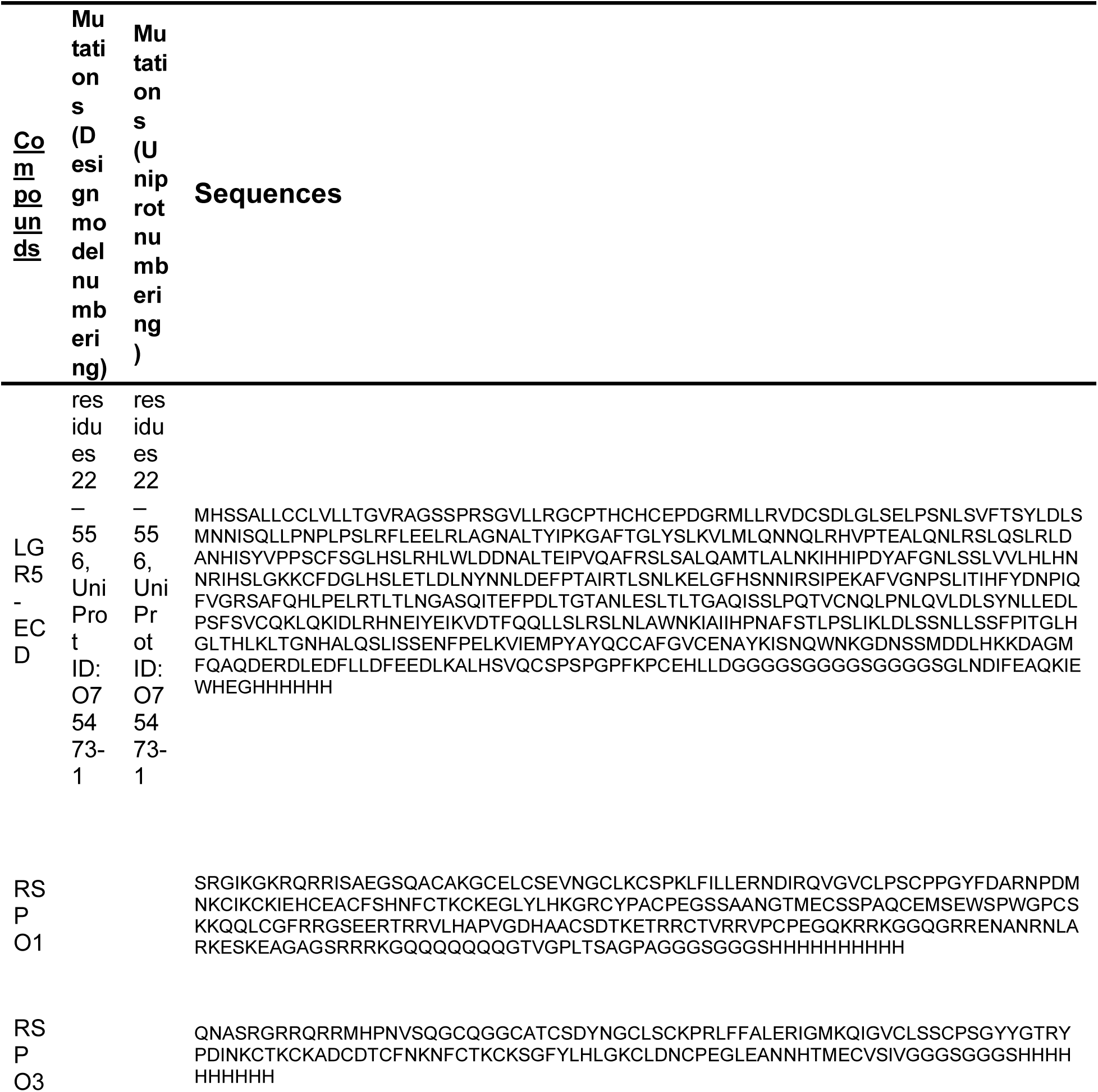
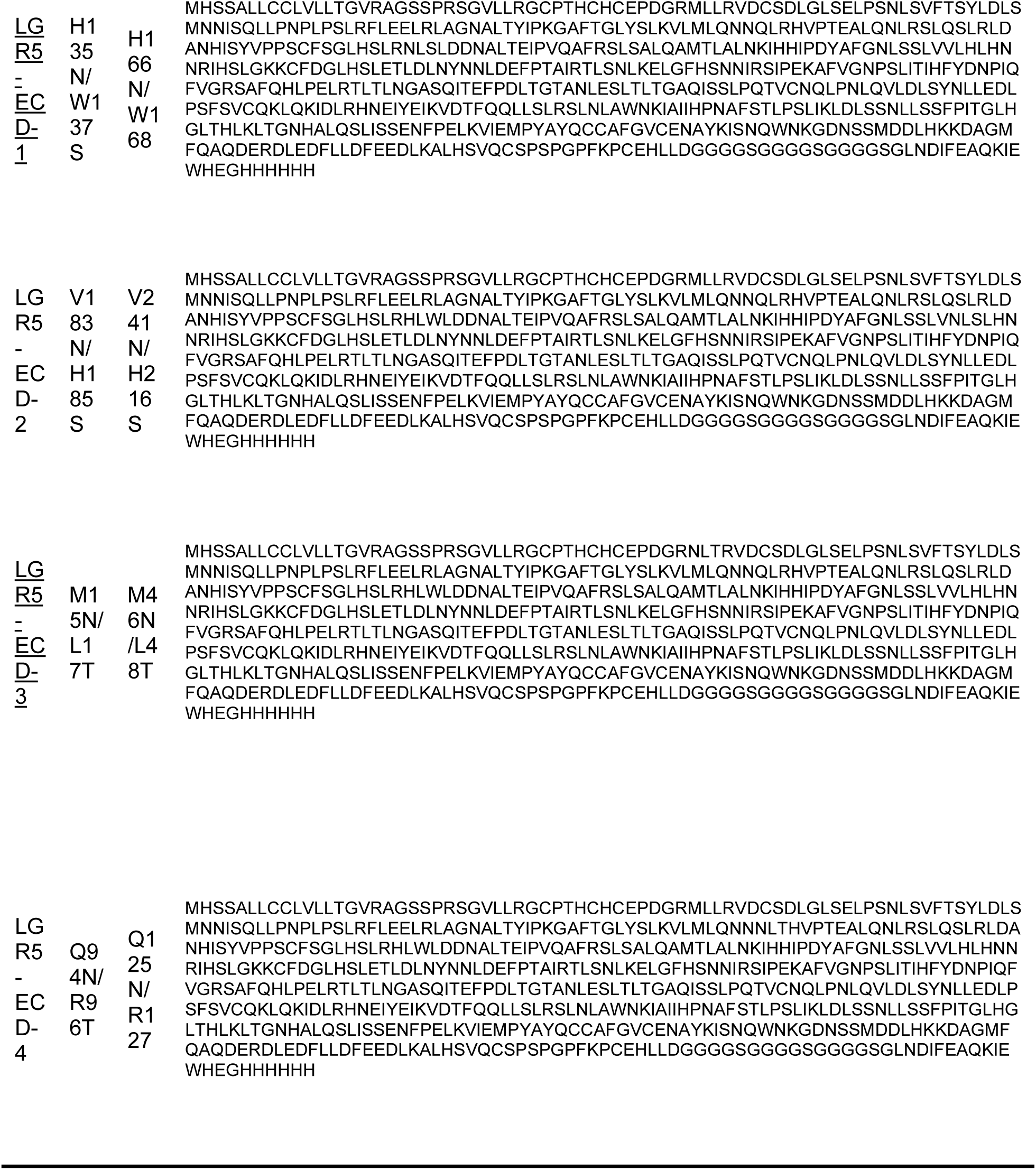
Biomaterials used in the research.

**Table S2:** Cryo-EM data collection, refinement, and validation statistics.

| Data collection and processing | Fab_LGR5-ECD_POC2-R3-23 |
| --- | --- |
| Microscope | Titan Krios G4 |
| Detector | Falcon 4 |
| Magnification | 96K |
| Voltage (kV) | 300 |
| Electron exposure (e-/Å²) | 50 |
| Defocus range (µm) | -1.0 -1.5 |
| Pixel size (Å) | 0.808 |
| Symmetry imposed | C1 |
| Initial particle images (no.) | 963,979 |
| Final particle images (no.) | 556,798 |
| Map resolution (Å) | 3.19 |
| FSC threshold | 0.143 |
| Refinement |  |
| Initial model used (PDB code) | Design VHH model |
| Map sharpening *B* factor (Å²) | -150.7 |
| Model composition |  |
| Protein residues | 808 |
| Ligands |  |
| B factors (Å²) |  |
| Protein Residues | 44.83 |
| Ligands |  |
| R.m.s. deviations |  |
| Bond lengths (Å) | 0.005 |
| Bond angles (°) | 0.656 |
| Validation |  |
| MolProbity Score | 1.71 |
| Clashscore | 7.54 |
| Poor rotamers (%) | 0.43 |
| Ramachandran plot |  |
| Favored (%) | 95.75 |
| Allowed (%) | 4.25 |
| Disallowed (%) | 0.00 |
| Model to map fit |  |
| CC_mask | 0.87 |
| CC_volume | 0.84 |

**Table S3:** Biophysical Characterization Summary.

| Design | K<sub>D</sub> (nM) | k<sub>a</sub> (1/Ms) | k<sub>d</sub> (1/s) | T<sub>m</sub>1 (°C) | Yield (mg/L) | SEC purity (%) |
| --- | --- | --- | --- | --- | --- | --- |
| R3-27 | 5.01E-07 | NA | NA | 54.2 | 450 | NA |
| R3-17 | 8.05E-07 | NA | NA | 66.8 | 594 | NA |
| R3-23 | 1.03E-07 | NA | NA | 74.3 | 543 | NA |

**Table S4:** Computational Filtering Metrics.

| Filter Stage | Designs Remaining | Metric Thresholds |
| --- | --- | --- |
| Raw Designs | 33168 | - |
| sc_value | 7557 | >0.6195 |
| cdrh3_dSASA_ratio | 6401 | >0.3188, <0.6965 |
| cdrh_dSASA_ratio | 4473 | >0.2862, <0.535 |
| boltz2 plddt | 4194 | >0.9 |
| boltz2 iptm | 2309 | >0,75 |
| chai1 iptm | 1763 | >0.25 |

### Supplementary Figures

**Figure S1.**
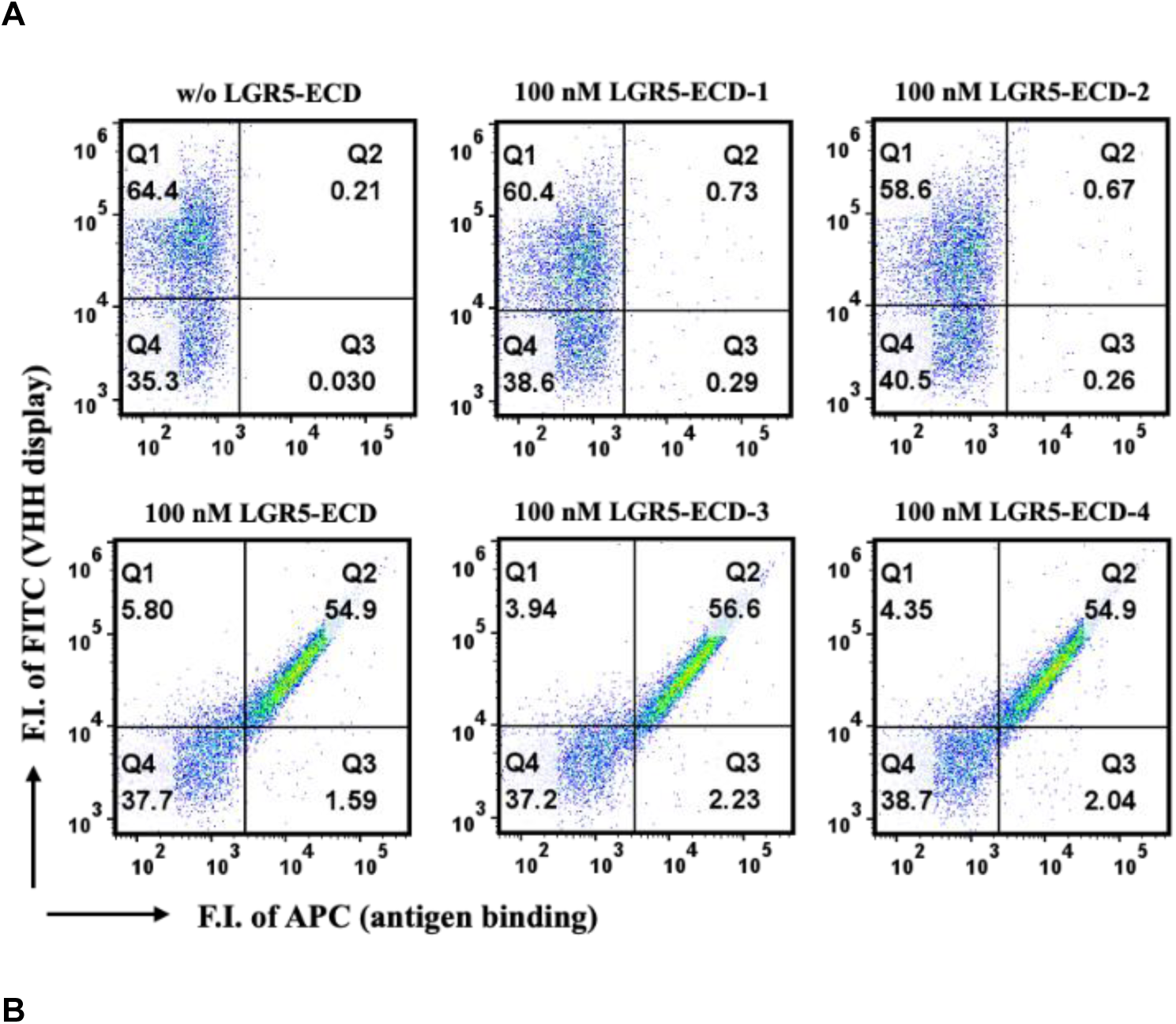
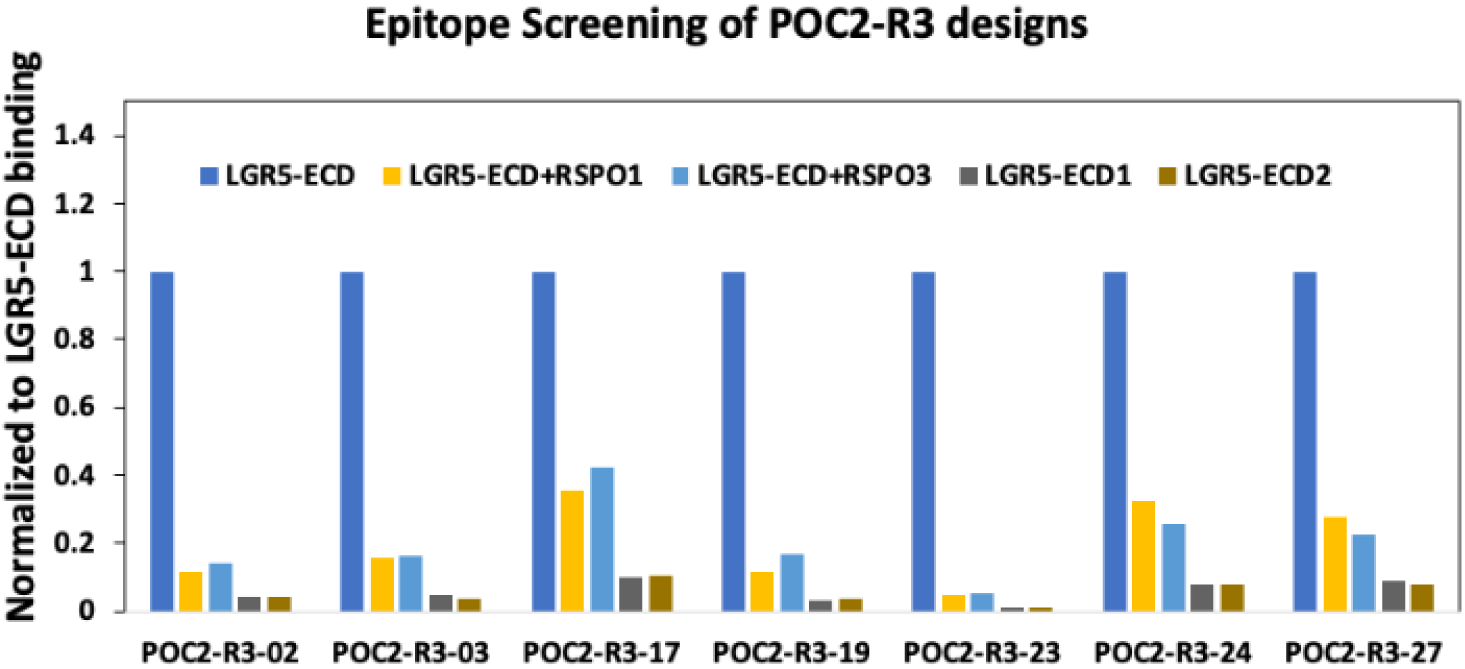
**FACS-based epitope characterization of *in silico* designs.** A) Yeast-displayed RSPO1 was incubated with 100 nM LGR5-ECD or its N-glycan variants. LGR5-ECD1 and LGR5-ECD2 share epitope overlap, while LGR5-ECD3 and LGR5-ECD4 serve as negative controls. The y-axis shows RSPO1 display efficiency (FITC fluorescence), and the x-axis shows RSPO1 binding to each compound (Avi-APC fluorescence). (B) Yeast-displayed in silico designs were incubated with 100 nM LGR5-ECD, its N-glycan variants, or 100 nM LGR5 plus 250 nM RSPO1/RSPO3. Display efficiency was measured using anti-FLAG-AF488, and binding efficiency using Avi-APC. Results were normalized to binding with LGR5-ECD.

**Figure S2.**
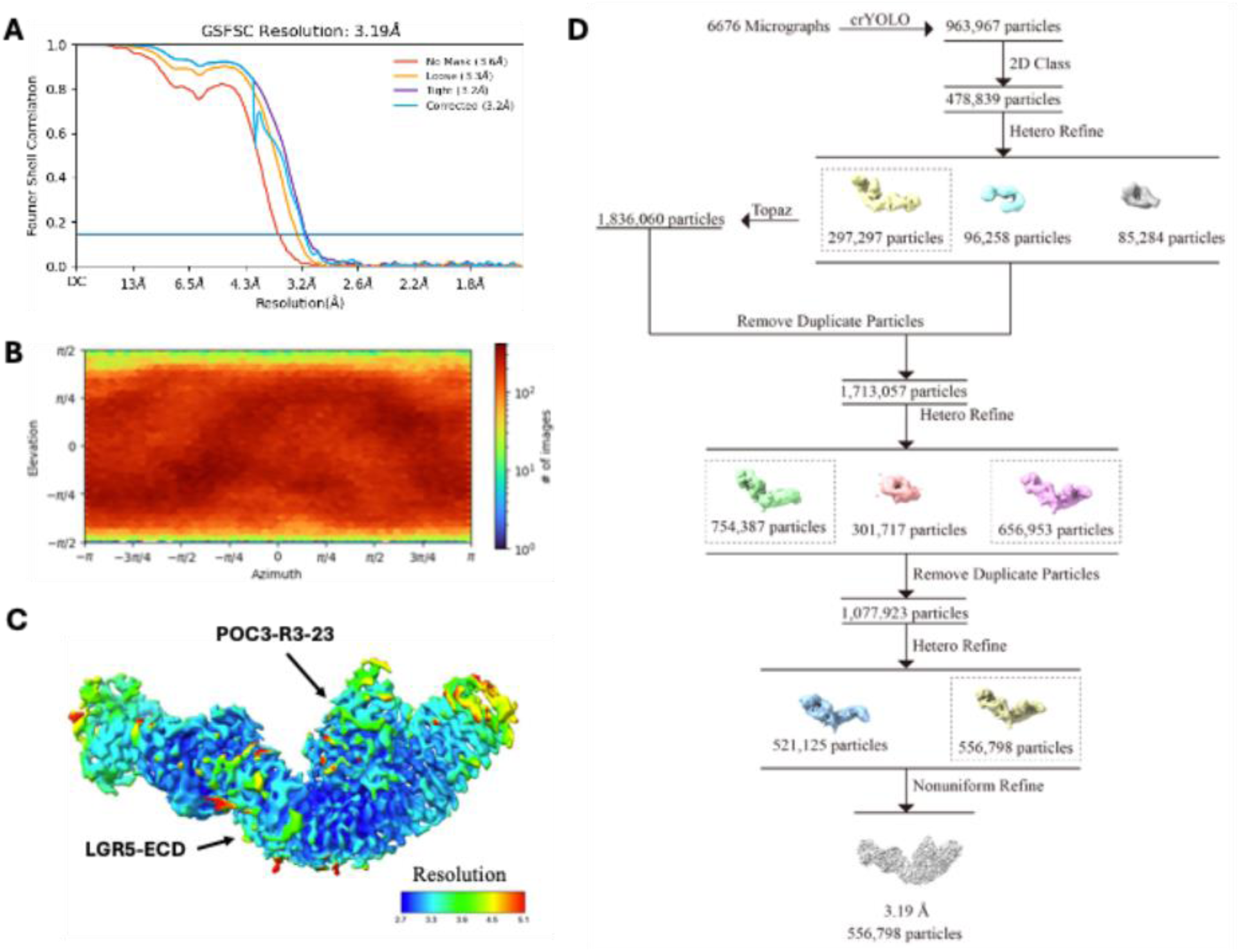
Cryo-EM complex structure data processing and statistics. (A) Global resolution plot showing final map resolution of 3.19 Å. (B) Orientational distribution confirming complete angular sampling. (C) Local resolution map (FSC = 0.143) with ∼3 Å resolution at the VHH–LGR5-ECD interface. (D) Cryo-EM processing workflow for the LGR5-ECD_POC2-R3-23 dataset. Refinement and filtering yielded a single binding pose during 3D classification, indicating high specificity of POC2-R3-23 for LGR5-ECD.

## References

1. Kaplon, H. & Reichert, J. M. Antibodies to watch in 2024. mAbs 16, 2297450 (2024).

2. Frenzel, A., Schirrmann, T. & Hust, M. Phage display-derived human antibodies in clinical development and therapy. mAbs 8, 1177–1194 (2016).

3. Muyldermans, S. Nanobodies: natural single-domain antibodies. Annu. Rev. Biochem. 82, 775–797 (2013).

4. Steeland, S., Vandenbroucke, R. E. & Libert, C. Nanobodies as therapeutics: big opportunities for small antibodies. Drug Discov. Today 21, 1076–1113 (2016).

5. De Genst, E., Silence, K., Decanniere, K., et al. Molecular basis for the preferential cleft recognition by dromedary heavy-chain antibodies. Proc. Natl. Acad. Sci. U.S.A. 103, 4586–4591 (2006).

6. Scully, M., Cataland, S. R., Peyvandi, F. et al. Caplacizumab treatment for acquired thrombotic thrombocytopenic purpura. N. Engl. J. Med. 380, 335–346 (2019).

7. Barker, N., van Es, J. H., Kuipers, J. et al. Identification of stem cells in small intestine and colon by marker gene Lgr5. Nature 449, 1003–1007 (2007).

8. de Sousa e Melo, F., Kurtova, A. V., Harnoss, J. M., et al. A distinct role for Lgr5⁺ stem cells in primary and metastatic colon cancer. Nature 543, 676–680 (2017).

9. Zimmermann, I., Egloff, P., Hutter, C. A. et al. Synthetic single-domain antibodies for the conformational trapping of membrane proteins. eLife 7, e34317 (2018).

10. Watson, J. L., Juergens, D., Bennett, N. R. et al. De novo design of protein structure and function with RFdiffusion. Nature 620, 1089–1100 (2023).

11. Dauparas, J., Anishchenko, I., Bennett, N. et al. Robust deep learning-based protein sequence design using ProteinMPNN. Science 378, 49–56 (2022).

12. Jumper, J., Evans, R., Pritzel, A. et al. Highly accurate protein structure prediction with AlphaFold. Nature 596, 583–589 (2021).

13. Cao, L., Coventry, B., Goreshnik, I. et al. Design of protein-binding proteins from the target structure alone. Nature 605, 551–560 (2022).

14. Jiang, L., Meng, R., Zhao, P. et al. De novo protein design enables the precise induction of RSV-neutralizing antibodies. Science 378, eabn0462 (2022).

15. Adolf-Bryfogle, J., Kalyuzhniy, O., Kubitz, M. et al. RosettaAntibodyDesign (RAbD): A general framework for computational antibody design. PLoS Comput. Biol. 14, e1006112 (2018).

16. Lau, J. L. & Roux, K. H. Computational design of antibody fragments. Curr. Opin. Struct. Biol. 83, 102708 (2023).

17. Greisen, P. Jr., Yi, L., Zhou, R., Zhou, J., Johansson, E., Dong, T., Liu, H., Johnsen, L. B., Lund, S., Svensson, L. A., Zhu, H., Thomas, N., Yang, Z. & Østergaard, H. Computational design of N-linked glycans for high-throughput epitope profiling. Protein Science 32, e4726 (2023).

18. Ellgaard, L. & Helenius, A. Quality control in the endoplasmic reticulum. Nat. Rev. Mol. Cell Biol. 4, 181–191 (2003).

19. Boder, E. T. & Wittrup, K. D. Yeast surface display for directed evolution. Nat. Biotechnol. 15, 553–557 (1997).

20. Leaver-Fay, A. et al. Rosetta3. Methods Enzymol. 487, 545–574 (2011).

21. Chothia, C. & Lesk, A. M. Canonical structures of immunoglobulins. J. Mol. Biol. 196, 901– 917 (1987).

22. de Lau, W. et al. The R-spondin/LGR5 module. Nature 478, 293–297 (2011).

23. Frank, C. et al. Scalable protein design using optimization in a relaxed sequence space. Science 386, 439–445 (2024).

24. Tyka, M. D. et al. Atomic-level protein energy landscape mapping. J. Mol. Biol. 405, 607– 618 (2011).

25. Arbabi-Ghahroudi, M. Camelid single-domain antibodies. Biochem. J. 474, 107–130 (2017).

26. Persson, H. et al. Glycan engineering for epitope occlusion. Nat. Commun. 4, 3034 (2013).

27. Zhou, Y. Arpeggia: A tool for geometric analysis of protein-protein interactions. Github (2024)

28. Fang, X. et al. A method for multiple-sequence-alignment-free protein structure prediction using a protein language model. Nat Mach Intell 5, 1087–1096 (2023).

29. Zheng, Z. et al. Structure-informed Language Models Are Protein Designers. Proceedings of the 40th International Conference on Machine Learning 202, 42317–42338 (2023).

30. Shusta, E. V., Kieke, M. C., Parke, E., Kranz, D. M. & Wittrup, K. D. Yeast polypeptide fusion surface display levels predict thermal stability and soluble secretion efficiency. J. Mol. Biol. 292, 949–956 (1999).

31. Deszyński, P., Młokosiewicz, J., Volanakis, A., Jaszczyszyn, I., Castellana, N., Bonissone, S., Ganesan, R. & Krawczyk, K. INDI—integrated nanobody database for immunoinformatics. Nucleic Acids Res. 50, D1273–D1281 (2022).

32. Wang, Y., Gong, X., Li, S., Yang, B., Sun, Y., Wang, Y., Shi, C., Yang, C., Li, H. & Song, L. xTrimoABFold: De novo antibody structure prediction without MSA. arXiv preprint arXiv:2212.00735 (2022).

33. James Dunbar, Konrad Krawczyk, Jinwoo Leem, Terry Baker, Angelika Fuchs, Guy Georges, Jiye Shi, Charlotte M. Deane, SAbDab: the structural antibody database, Nucleic Acids Research, Volume 42, Issue D1, 1 January 2014, Pages D1140–D1146

